# Female Assamese macaques bias their affiliation to paternal and maternal kin

**DOI:** 10.1101/714253

**Authors:** Delphine De Moor, Christian Roos, Julia Ostner, Oliver Schülke

**Affiliations:** Department of Behavioural Ecology, University of Goettingen, Germany; Primate Genetics Laboratory, German Primate Center, Leibniz Institute for Primate Research, Goettingen, Germany; Leibniz-ScienceCampus Primate Cognition, Goettingen, Germany; Research Group Primate Social Evolution, German Primate Center Leibniz Institute for Primate Research, Goettingen, Germany

**Author notes:** Contributed equally as senior authors (Julia Ostner, Oliver Schülke). Corresponding author: Delphine De Moor Kellnerweg 6, 37077 Göttingen, Germany.

**Keywords:** kin discrimination, kin selection, nepotism, relatedness, social bonds

## Abstract

Forming strong social bonds leads to higher reproductive success, increased longevity and/or increased infant survival in several mammal species. Given these adaptive benefits, understanding what determines partner preferences in social bonding is important. Maternal relatedness strongly predicts partner preference across many mammalian taxa. Although paternal and maternal kin share the same number of genes, and theoretically similar preferences would therefore be expected for paternal kin, the role of paternal relatedness has received relatively little attention. Here, we investigate the role of maternal and paternal relatedness for female bonding in Assamese macaques (*Macaca assamensis*), a species characterized by a relatively low male reproductive skew. We studied a wild population under natural conditions using extensive behavioural data and relatedness analyses based on pedigree reconstruction. We found stronger social bonds and more time spent grooming between maternal kin and paternal half-sisters compared to non-kin, with no preference of maternal over paternal kin. Paternally related and non-related dyads did not form stronger bonds when they had less close maternal kin available, however we would need a bigger sample size to confirm this. As expected given the low reproductive skew, bonds between paternal half-sisters closer in age were not stronger than between paternal half-sisters with larger age differences, suggesting that age similarity was not the mechanism by which paternally related individuals recognized each other. An alternative way through which paternal kin could get familiarized is mother- and/or father-mediated familiarity.

## INTRODUCTION

Many group-living animals show considerable variation in how they interact with other group members, forming strong relationships with only a few of their group mates and weak relationships with the rest of the group (Cameron, Setsaas, & Linklater, 2009; Diaz-Aguirre, Parra, Passadore, & Möller, 2018; Silk, Altmann, & Alberts, 2006; J. E. Smith et al., 2010). As gregarious animals repeatedly associate with each other, the quality of their interactions can affect subsequent interactions, reinforcing the nature of their relationships (Hinde, 1976). Over time, such reinforcement leads to stable partner preferences in affiliation or social bonds (Silk, Cheney, & Seyfarth, 2013). Investing time and energy in the formation and maintenance of these bonds might be adaptive in terms of inclusive fitness benefits, mutual benefits, or reciprocal exchanges (Clutton-Brock, 2009; Kummer, 1978). In several mammal species, individuals forming strong stable social bonds have a higher reproductive success, increased longevity and/or increased infant survival compared to more weakly bonded group mates (Brent, Chang, Gariépy, & Platt, 2014; Ostner & Schülke, 2018; but see Kalbitzer et al., 2017; Thompson & Cords, 2018). Given this strong selective pressure, it is crucial to understand what factors underlie partner preferences in social bonding.

An important factor shaping adult partner preferences across most mammalian taxa is maternal relatedness (Archie, Moss, & Alberts, 2006; Berman, 2015; Möller, 2012; Rossiter, Jones, Ransome, & Barratt, 2002; Seyfarth & Cheney, 2012; Silk, 2009; J. E. Smith, 2014; J. E. Smith et al., 2010). Because of the extended maternal care that is characteristic of most mammals, infants are closely spatially associated with their mother, which may promote the formation of strong social bonds that are maintained into adulthood (Broad, Curley, & Keverne, 2006). Females of many mammal species are philopatric; they spend their entire life in their natal group, surrounded by their maternal kin (Greenwood, 1980). In these species, mothers are imbedded into a social network of their own close maternal kin, and the strong mother-offspring bond in turn facilitates the development of bonds between close maternal kin, as infants are indirectly familiarized to their maternal siblings, aunts and grand-mother very early on (i.e. mother-mediated familiarity; Widdig, 2007). Maternal kin are readily available and easily recognizable, which could facilitate tolerance, affiliation and cooperation among maternally related females (Cords & Nikitopoulos, 2015; Silk, 2009). More ultimately, a kin bias in affiliation towards maternal kin could arise due to the indirect fitness benefits individuals gain from cooperating with related individuals, according to kin selection theory (Hamilton, 1964).

Given that kin selection is blind to the origin of shared genes (Hamilton, 1964), one would expect a similar bias in affiliation towards paternal kin as towards maternal kin. A key issue, however, is paternal kin recognition (Strier, 2004; Widdig, 2007). In many mammal species, females mate with more than one male during their fertile period (Jennions & Petrie, 2000). Paternity is concealed in those species, which might hamper paternal kin recognition and, by extension, the development of a bias towards paternal kin. Still, several mechanisms of paternal kin recognition have been put forward. In species with a high male reproductive skew and a short alpha male tenure relative to the interbirth interval, infants that are born close in time are likely sired by the alpha male and thus have a high chance of sharing the same father. As close-aged individuals grow up together, age proximity could be a potential mechanism through which paternal kin are familiarized to each other (i.e. familiarity via age proximity; Altmann, 1979; Widdig, 2013). Alternatively, paternal kin could recognize each other through mother-or father-mediated familiarity (Widdig, 2007). Mothers may mediate familiarity between paternal siblings directly, or infants could get familiarized through their common father if mothers affiliate with the father after birth. Similarly, fathers could mediate familiarity if they affiliate with the mother, or if they provide paternal care to their offspring. Independent of familiarity, individuals could recognize their kin using phenotypic cues such as appearance, odour and vocalizations (i.e. phenotypic matching; Mateo, 2017; Widdig, 2007).

Primates have been particularly well studied on kin biases in sociality, because they live in stable, complex groups often containing both paternal and maternal kin, as well as non-related individuals (Chapais & Berman, 2004; Silk et al., 2006). While maternal relatedness is generally a strong predictor of bond formation (Silk, 2009; but see Candiotti et al., 2015), research on paternal kin biases is still limited and the results are mixed (Berman, 2015). So far, paternal kin biases in primates have only been reported in cercopithecines (Charpentier et al., 2012; Cords, Minich, Roberts, & Sleator, 2018; Lynch, Di Fiore, Lynch, & Palombit, 2017; Silk et al., 2006; Widdig, Nurnberg, Krawczak, Streich, & Bercovitch, 2001), while no biases towards paternal kin have been found in other primate taxa (Langergraber, Mitani, & Vigilant, 2007; Perry, Manson, Muniz, Gros-Louis, & Vigilant, 2008; Wikberg, Ting, & Sicotte, 2014).

Paternal kin biases are usually less pronounced than biases towards maternal kin (Charpentier et al., 2012; Lynch et al., 2017; Silk et al., 2006; Widdig et al., 2001). This might reflect the lower certainty of recognizing paternal kin compared to maternal kin (Berman, 2015). Additionally, the presence and strength of paternal kin biases evidently depend on factors such as demographic and mating patterns (Strier, 2004), like the availability of both maternal and paternal kin as potential bonding partners. For example, in two studies on the same population of yellow baboons (*Papio cynocephalus*), paternal half-sisters were preferred to the same extent as maternal half-sisters in a period when relatively few females had close maternal kin available, while social bonds between paternal half-sisters were of intermediate strength between maternal kin and non-kin in a later period when more females had close maternal kin available (Silk et al., 2006; K. Smith, Alberts, & Altmann, 2003). Individuals might thus form close bonds with paternal kin to compensate for the absence of the preferred close maternal kin (Silk et al., 2006).

Male reproductive skew is another major influence on paternal kin biases, determining the availability of paternal kin and the likelihood for individuals born close in time to be paternally related and thus the reliability of age proximity as a proxy for paternal relatedness (Altmann, 1979). It should be noted that familiarization with paternal kin via age similarity has been questioned on theoretical grounds (Langergraber, 2012). While seasonal reproduction should favour familiarization via age similarity because infants grow up together, increased seasonality also decreases monopolization potential and thus reproductive skew among males, reducing the chance that age mates are paternal kin. Kin biases towards paternal kin are stronger within than between age cohorts in some species (yellow baboons: Silk et al., 2006; rhesus macaques - *Macaca mulatta*: Widdig et al., 2001) but not in others, even though paternity concentration in the alpha male is rather high (mandrills - *Mandrillus sphinx*,, alpha male skew: 76.2%; Charpentier, Peignot, Hossaert-McKey, & Wickings, 2007; blue monkeys - *Cercopithecus mitis*, resident male skew: 61%; Cords et al., 2018).

Here, we study kin biases in a population of Assamese macaques (*Macaca assamensis*) under natural conditions, combining longitudinal behavioural data with genetic analyses of relatedness based on pedigree reconstruction. Assamese macaques have a relatively low reproductive skew (29% alpha male paternity: Sukmak, Wajjwalku, Ostner, & Schülke, 2014), and a relatively long alpha male tenure relative to the interbirth interval, with males often being able to keep their alpha position for consecutive mating seasons (Ostner, Vigilant, Bhagavatula, Franz, & Schülke, 2013). Familiarization of paternal kin through age proximity is thus unlikely. Still, because of the long male tenure, many individuals might have paternal kin available in the group, necessitating even more so to distinguish the few paternal kin from the many non-kin. We predict (1) that, like in several other cercopithecine and mammal species, adult females bias their affiliation towards maternal kin, (2) that affiliation is also biased to paternal kin, with intermediate bond strength between maternal kin and non-kin, (3) that age proximity does not affect bond strength between paternal kin, and (4) that paternal kin form stronger social bonds when either one or both of the partners have few close maternal kin available, to compensate for the lack of preferred maternally related bonding partners.

## MATERIAL AND METHODS

### Study site and study population

We studied a population of fully habituated Assamese macaques ranging in their natural habitat at the Phu Khieo Wildlife Sanctuary in north-eastern Thailand (PKWS, 16°05–35’N & 101°20– 55’E). Part of the more than 6500 km^2^ interconnected and well-protected Western Isaan forest complex, the sanctuary covers an area of >1600 km^2^, with elevations ranging from 300 to 1300m above sea level (Schülke, Pesek, Whitman, & Ostner, 2011). The pristine forest is composed of dense, dry evergreen vegetation with patches of bamboo forest and harbours a diverse community of large mammals and predators (Borries, Larney, Kreetiyutanont, & Koenig, 2002).

This study is part of a long-term research project on this population of Assamese macaques. All individuals could be reliably identified based on natural variation in appearance. We collected data on one group in 2007/2008 and 2010/2011, on two groups in 2013/2014, and on four groups in 2015/2016 and 2016/2017. Each study period captured at least one year, including the mating season from October to February and the non-mating season from March to September (Fürtbauer, Schülke, Heistermann, & Ostner, 2010). Assamese macaques are characterized by female philopatry and male dispersal. At any time, all groups included several adult males (7.4 (mean) ± 3.6 (SD)) and adult females (11.8 (mean) ± 3.9 (SD)), as well as a large number of immatures. This study focused on adult females (n=59), with females defined as adult in the mating season of their first conception. For 25 of these females the mother and exact age are known from direct observation. The other 34 females were already present in the population when the project started in 2006, so no information on their mother was available and age was estimated based on morphology and behaviour.

This study was conducted completely non-invasively and adhered to the ASAB/ABS Guidelines for the Use of Animals in Research. The Department of National Parks, Wildlife and Plant Conservation (DNP) and the National Research Council of Thailand (NRCT) authorized data collection and export of samples with a benefit sharing agreement (permit numbers: 0004.3/3618, 0002.3/2647, 0002/17, 0002/2424, 0002/470).

### Behavioural data collection

We followed the study groups from dawn to dusk, from sleep tree to sleep tree. During 40 minutes continuous focal animal observations, we recorded frequencies and durations along with actor and receiver of affiliative behaviours (grooming, body contact and approaches into and departures from a 1.5 m radius around the focal individual). An effort was made to balance focal animal protocols across individuals and time of day per study period (143 (mean) ± 13 (SE) hours/female, 8412 hours total).

### Composite sociality index

To measure the strength of affiliative relationships among adult females, we used a modified dyadic composite sociality index (CSI; Silk et al., 2006). In this study, the CSI is based on the frequency (F) and duration (D) of the following social behaviours: proximity within 1.5m (P), body contact (B) and grooming (G). It is calculated as the sum of the dyadic frequency and duration of every behaviour, divided by the mean group frequency or duration of that behaviour and then divided by the number of behaviours (=6): 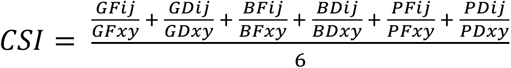, with GFij = grooming frequency for dyad ij, GFxy = mean grooming frequency for all dyads in group x and year y, etc. CSI values thus express the extent to which a dyad’s affiliation deviates from the average female dyad in a given group and period. The CSI ranges from 0 to infinity, with an average of 1 and stronger than average relationships for the respective group and period indicated by values higher than 1. Time spent in proximity was subtracted from the time spent in body contact and time spent in body contact was subtracted from time spent grooming to not double the impact of nested interactions. All behaviours were corrected for dyadic observation time (i.e. the sum of observation time for each dyad member during which the other dyad member was adult and present as well). Before calculating CSI values, we checked that mean frequencies of each behaviour exceeded 2, to ensure that enough data was available to compute a meaningful CSI value for a given period. All six behavioural measures included in the index were significantly positively correlated (row-wise matrix correlations for all pair-wise combinations: average row-wise tau values ranging from 0.34 to 0.85 across years and groups).

### Faecal sample collection

Faecal samples for genetic analyses had been collected non-invasively since the start of the project in 2006. About 5g of material was scratched from the surface of excrements immediately after defecation by an identified individual. We applied the two-step storage procedure (Nsubuga et al., 2004): the faecal material was stored in a 50mL falcon tube (62.559.001, Sarstedt, Nümbrecht, North Rhine-Westphalia, Germany) containing 30ml of 97% ethanol and labelled with animal ID, time, date and collector. Samples were collected opportunistically throughout the day. Twenty-four to 36 hours after collection, the ethanol was carefully poured off, and the remaining solid material was transferred to a new 50ml falcon tube containing silica gel beads (112926-00-2, Intereducation Supplies Co., Ltd., Bangkok, Thailand). Samples were stored at room temperature in the dark until shipped to the German Primate Center (DPZ) for further analysis. Multiple independent faecal samples have been collected for each individual since 2006 and are stored at −20°C until DNA extraction.

### DNA extraction & genotyping

DNA was extracted from 100-150mg of faeces at the Primate Genetics Laboratory (DPZ, Goettingen), using the First-DNA all-tissue kit (D1002000, GEN-IAL GmbH, Troisdorf, North Rhine Westphalia, Germany). We followed the manufacturer’s protocol for DNA extraction from faeces, using DNA LoBind Tubes (0030108051, 0030108078, Eppendorf AG, Hamburg, Germany) in all steps with a final resuspension of DNA in 50µl HPLC (high-performance liquid chromatography) water. We measured the DNA concentration of the extracts using NanoDrop (Spectrophotometer ND-1000, PEQLAB Biotechnologie GmbH, Erlangen, Bavaria, Germany), and diluted them to a concentration of 100 ng/µl for further use in standard PCR.

Eighty-eight individuals of the same population had been genotyped previously at 15 autosomal microsatellite loci (Table 1; Sukmak et al., 2014). We continued this effort for the remaining 48 individuals of the population that were adult before 2016, using the same procedures, and included 2 additional microsatellite loci for all 136 individuals (D1s548 & D7s2204; established for crested macaques -*Macaca nigra* in Engelhardt, Muniz, Perwitasari-Farajallah, & Widdig, 2017). We followed the two-step multiplex PCR approach (Arandjelovic et al., 2009): first, we used 1µl of DNA extract (100ng/µl dilution) and all primer pairs (10mM) in the multiplex reaction, then diluted the PCR product 1:1000 and used 5µl of that dilution and one primer pair (10mM) per reaction for the singleplex reactions. We added BSA (bovine serum albumin, 20mg/ml) and MgCl_2_ (25 mM) to all reactions to enhance PCR performance and used SuperTaq DNA polymerase (020503, Cambio Ltd, Cambridge, United Kingdom) combined with TaqStart Antibody (639250, Takara Bio Europe, Saint-Germain-en-Laye, France) to prevent non-specific amplification. Multiplex reaction conditions for all primer pairs were initial denaturation at 94 °C for 9 min, 29 cycles of denaturation at 94 °C for 20 s, annealing at 50 °C for 30 s, and extension at 72 °C for 30 s, then a final extension at 72 °C for 4 min. Singleplex reactions followed the same conditions, but with an annealing temperature of 51.9 °C. For the singleplex reactions, the forward primers were fluorescently labelled at the 5’ end (Table 1). After running the singleplex products together with a non-template control on 1.5% agarose gels, we made dilutions (pure product to 1:50 dilution) for every sample based on the brightness of the band. We ran these dilutions on an ABI 3130xL sequencer (Applied Biosystems) and used the software Geneious (version 11.0.4, www.geneious.com) to determine product sizes relative to the ROX HD400 internal size standard.

**Table 1:**
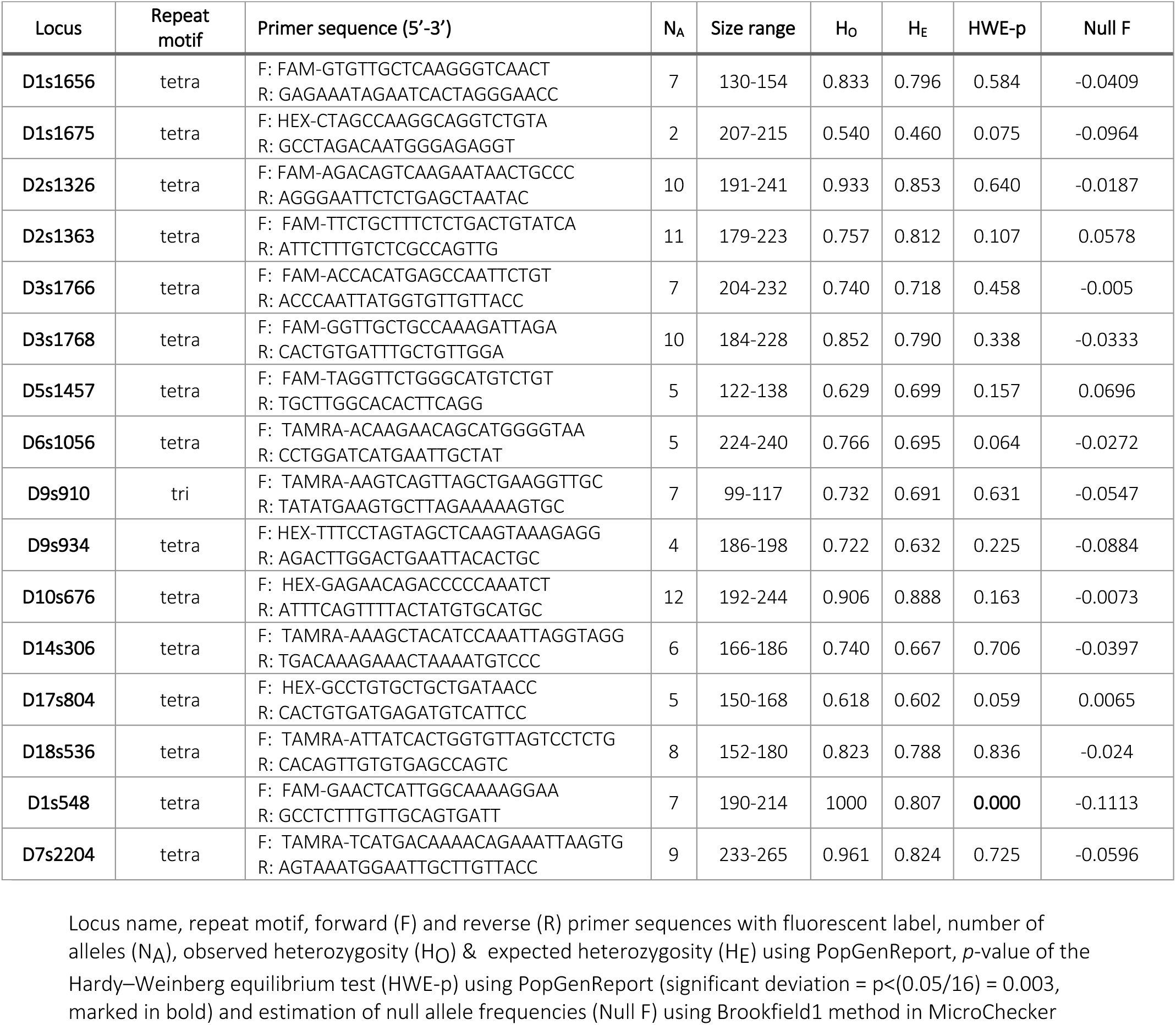
Information on the 16 microsatellite loci used in this study

To control for possible misidentification of individuals in the field, we genotyped all individuals from two different samples; to control for mistakes in the lab, we genotyped every extraction two times independently (multiple-tube approach; Navidi, Arnheim, & Waterman, 1992; Taberlet et al., 1996), leading to four genotypes per individual. An allele was accepted when it appeared at least in three of those four genotypes, and at least once for every independent extract. To protect against false homozygosity from allelic dropout, we accepted a locus as homozygous only if the allele occurred independently at least seven times, according to guidelines for attaining 99% statistical confidence in genotyping (Taberlet et al., 1996). One locus (D8s1106) was excluded as genotyping often failed, and a closer look at the sequence of the forward primer binding region showed that an insertion in the macaque DNA sequence (compared to human DNA for which the primer was developed) caused the primer to not bind properly. We checked the genotypes with Micro-Checker (version 2.2.3; Van Oosterhout, Hutchinson, Wills, & Shipley, 2004) and found no evidence for the occurrence of null alleles, large allele drop-out or scoring error due to stutter. For 133 of the 136 individuals, genotypes were generated for at least 8 of the 16 remaining loci (typed at 13.2 (mean) ± 1.6 (SD) loci, mean proportion of loci typed 81.66%). The 3 individuals with <8 loci typed were excluded from parentage analyses. We tested that all loci were in Hardy-Weinberg equilibrium using PopGenReport (version 3.0.0; Adamack & Gruber, 2014), and found that locus D1s548 departs from Hardy-Weinberg equilibrium, which is attributable to the presence of close relatives in the study group. Mean number of alleles per locus was 7.2, mean polymorphic information content (PIC) was 0.69 and mean expected heterozygosity was 0.74, all calculated using Cervus (version 3.0.0; Kalinowski, Taper, & Marshall, 2007).

Apart from the 16 microsatellite loci, we also genotyped all individuals at the hypervariable region I of the mitochondrial D-loop region. Mitochondrial DNA (mtDNA) is maternally inherited, so candidate mothers for parentage analysis (see below) were restricted to females with a matching mtDNA haplotype, which helped reduce misclassification rates (Kopps, Kang, Sherwin, & Palsboll, 2015). PCRs were performed using the primers 5’-AAATGAACCTGCCCTTGTAGT-3’ and 5’-GAGGATAGAACCAGATGTCC-3’ with initial denaturation at 94 °C for 5 min, 40 cycles of denaturation at 94 °C for 1 min, annealing at 50 °C for 1 min, and extension at 72 °C for 1.5 min, then a final extension at 72 °C for 5 min. PCR products together with a non-template control were checked on 1% agarose gels and then cleaned with the Qiagen PCR Purification Kit and sequenced using an ABI 3130xL sequencer and the BigDye Cycle Sequencing Kit (Applied Biosystems). In total we found four mtDNA haplotypes: one was represented only in two of the four observed groups, two others only in the other two observed groups (that were originally one group) and one was only found in immigrant males.

### Parentage assignment

For every individual a conservative set of candidate parents was determined. All males that were at least five years old at the time of birth of the individual were included as potential fathers, all females that had the same mtDNA haplotype as the individual and were at least five years old at the time of birth of the individual were included as potential mothers. From those candidate parents, we assigned parentage using both the pairwise likelihood-basis paternity analysis of Cervus (version 3.0.0; Kalinowski et al., 2007) and the full-pedigree likelihood method of Colony (version 11.3.0; Jones & Wang, 2010). Cervus assigns a parent by calculating the difference in log-likelihood ratios (LOD score, the likelihood of parentage of a particular parent compared to the likelihood of parentage of an arbitrary parent; Kalinowski et al., 2007). Colony on the other hand considers likelihoods over the full pedigree, comparing different clusterings of individuals, where individuals in a cluster are related via sibship or shared parentage. Cervus assigned more parents than did Colony, but with the exception of one sire, both Cervus and Colony assigned the same parents to individuals, confirming the reliability of our parentage assignments. We therefore included all parents assigned at a strict (95% likelihood) level by Cervus in the analysis. Relatedness values (r) were also calculated for every dyad, by using the R package Related (version 1.0; Pew, Muir, Wang, & Frasier, 2015) and the triadic likelihood estimator (trioML; Wang, 2007).

Based on the parentage analyses, dyads were classified as mother-daughters, maternal half-sisters, paternal half-sisters and non-kin. Individuals who belonged to more than one kin category, such as full siblings, were excluded from the analysis. Non-kin dyads were defined as dyads for whom both parents of each dyad member were known and not shared, and had a relatedness value lower than 0.125. Out of the 513 female-female dyads for which we had behavioural data (i.e. who were adults living in the same group at some point during the 6 years of observational data available), we were able to assign a total of 161 to one of those kin classes (Table 2). The remaining dyads either belonged to more than one kin category (3 full sibling dyads, 2 mother-offspring dyads who were also paternal half-sisters), shared no parents but did not fit the threshold of r < 0.125 for non-kin (130 dyad) or we lacked parentage information to reliably assign them to a kin class (217 dyads).

**Table 2:** Number of adult females assigned to each kin category and average relatedness (TrioML after Wang, 2007) of all dyads in that kin class

| Kin class | Non-kin | Paternal half-sisters | Maternal half-sisters | Mother-daughters |
| --- | --- | --- | --- | --- |
| Nr. of dyads | 94 | 12 | 21 | 34 |
| Average relatedness | 0.033 | 0.285 | 0.244 | 0.472 |

### Statistical analyses

All analyses were run in R (version 3.4.4; R Development Core Team, 2018), using the Rstudio interface (version 1.0.153; RStudio Team, 2016). Since our data set was longitudinal, with repeated measures across several years and groups we ran linear mixed models, to allow to control for non-independence of the data, using the R package lme4 (Bates, Mächler, Bolker, & Walker, 2015). For all models, we checked the assumption of normally distributed and homogeneous residuals by visually inspecting a qqplot and scatterplot of the residuals plotted against the fitted values. None of the models showed obvious deviations from these assumptions. Model stability was tested by excluding data points one by one from the data and comparing the model estimates for the subsets to the estimates for the full data set. No influential cases were found in any of our models. We checked that the predictors of interest were not collinear by calculating Variance Inflation Factors (VIF) using the function vif of the R package car (Fox & Weisberg, 2011) applied to a standard linear model excluding the random effects. VIFs were close to one for all models, indicating collinearity not to be an issue. Covariates were z-transformed to a mean of 0 and standard deviation of 1 to allow for comparison of effect sizes. All statistical analyses were performed on a dyadic level. To make sure that assigning a member of the dyad to ID1 or ID2 had no influence, the models were iterated 10,000 times, with random assignment of dyad members to either ID1 or ID2. For all models, we performed a full-null model comparison, with the null model containing only the control predictors, to test whether inclusion of predictors improved model fit. P-values for significance of each level of the categorical predictors were obtained by using the function glht or the R package multicomp, that corrects for multiple comparisons (Hothorn, Bretz, & Westfall, 2008).

#### Models 1&2: social bond strength and time spent grooming in the different kin classes

To analyse whether kin class influences dyadic social bond strength, we ran a Linear Mixed Model with kin class as the predictor, and the square root of the dyadic CSI value as the response. Group, year, dyad, ID1 and ID2 were included as random effects, and kin class within group and within year were included as random slopes. We ran the model once with non-kin as the reference level, to test whether the three related kin classes (maternal and paternal half-sisters and mother-daughters) differed from non-kin in their social bond strength, and then releveled the variable kin class to paternal half-sisters, to test for differences between paternal half-sisters and close maternal kin. To check the robustness of the results we ran the same model with the square root of time spent grooming – an unambiguously friendly and directed behaviour – as the response. For both these models all assigned dyads were included: 94 non-kin dyads (171 observations), 12 paternal half-sisters (21 observations), 21 maternal half-sisters (54 observations) and 34 mother-daughter dyads (73 observations).

#### Model 3: the effect of close maternal kin availability on paternal kin biases

To assess whether availability of maternal kin influences bond strength with paternal half-sisters compared to non-kin, we ran a Linear Mixed Model with the sum of the number of close maternal kin (i.e. adult and alive mother, daughters and maternal half-sisters) for both partners of each dyad and kin class as predictor variables, and the square root of the dyadic CSI value as the response. Dyad, ID1 and ID2 were included as random effects. We restricted this analysis to two of the four observed groups on which we have data since 2006, as we have sampled more mothers of the currently adult females in those groups and the proportion of assigned dyads is higher (49%). Individuals had on average 1.97 close maternal kin available (ranging from 0 to 5), the sum of close maternal kin available to both partners of a dyad was on average 3.94 (ranging from 1 to 8). For this model we had 81 non-kin dyads (146 observations) and 9 paternal half-sisters (16 observations).

#### Models 4&5: effect of age difference on paternal kin bias

To test whether the bias towards paternal kin was more pronounced for paternal half-sisters closer in age, we ran a Linear Mixed Model with the square root of the age difference, kin class and the interaction between age difference and kin class as predictor variables, and the square root of the dyadic CSI value as the response. Group, year, dyad, ID1 and ID2 were included as random effects, but no random slopes were included in this model, as there were not enough levels for kin class or values for age difference with at least two dyads per group or year. Age difference ranged from 0 to 11 years. We also ran the same model with a binary predictor same age cohort or not (i.e. born in the same year or not) instead of age difference (21 dyads born in the same cohort, 36 observations; 85 dyads born in a different cohort, 156 observations). For both these models we used a subset of the data including only non-kin (94 dyads, 171 observations) and paternal half-sisters (12 dyads, 21 observations).

## RESULTS

### Social bond strength and time spent grooming in the different kin classes (Models 1&2)

Overall, the full model with social bond strength as the response and including kin class as predictor was highly significant compared to the null model (likelihood ratio test: χ^2^ = 14.59 [12.03, 16.25], df = 3, p = 0.002 [0.001, 0.007], mean [minimum, maximum] over 10,000 tests), meaning that the multi-level variable kin class had a significant effect on the strength of dyadic social bonds (reference level: non-kin; marginal R^2^= 0.14 [0.12, 0.16], mean [minimum, maximum] over 10,000 tests). More specifically, we found significantly stronger bonds between dyads of all related kin classes (paternal half-sisters, maternal half-sisters and mother-daughters) compared to non-kin dyads (Figure 1 & Table 3). After releveling the variable kin class to paternal half-sisters, we did not find significant differences in social bond strength between paternal half-sisters and maternal kin. The full model with time spent grooming as the response and including kin class as predictor was highly significant compared to the null model as well (likelihood ratio test: χ^2^ = 12.71 [11.50, 14.87], df = 3, p = 0.005 [0.002, 0.009], mean [minimum, maximum] over 10,000 tests), meaning that the multi-level variable kin class also had a significant effect on the time spent grooming (reference level: non-kin; marginal R^2^= 0.18 [0.16, 0.20], mean [minimum, maximum] over 10,000 tests). (Figure 2 & Table 4).

**Table 3:** Effect of kin class on dyadic social bond strength

| term | Estimate | Standard Error (SE) | t score | p value |
| --- | --- | --- | --- | --- |
| intercept | 0.825 [0.797, 0.860] | 0.052 [0.047, 0.063] | - | - |
| paternal half-sisters | 0.311 [0.212, 0.373] | 0.117 [0.109, 0.120] | 2.665 [1.832, 3.229] | 0.03 [0.03, 0.04] |
| maternal half-sisters | 0.232 [0.157, 0.306] | 0.082 [0.078, 0.091] | 2.813 [1.951, 3.648] | 0.01 [0.01, 0.01] |
| mother-daughter | 0.502 [0.461, 0.541] | 0.125 [0.112, 0.138] | 3.992 [3.457, 4.588] | 0.0001 [0.0001, 0.0005] |

model estimates (mean [minimum, maximum]) for the 10,000 iterations of the model with non-kin as reference level linear mixed model: $\text{sqrt(CSI)} \sim \text{kin class} + (1|\text{kin class}||\text{group}) + (1|\text{kin class}||\text{year}) + (1|\text{dyad}) + (1|\text{focal\_animal}) + (1|\text{action\_partner})$ t score & p value for intercept not shown because of limited interpretation

**Table 4:** Effect of kin class on time spent grooming in a dyad

| term | estimate | Standard Error (SE) | t score | p value |
| --- | --- | --- | --- | --- |
| intercept | -1.628 [-1.706, -1.567] | 0.102 [0.092, 0.122] | - | - |
| paternal half-sisters | 0.464 [0.380, 0.539] | 0.180 [0.176, 0.182] | 2.580 [2.131, 3.013] | 0.03 [0.03, 0.04] |
| maternal half-sisters | 0.370 [0.295, 0.500] | 0.124 [0.122, 0.184] | 2.983 [2.132, 3.389] | 0.009 [0.008, 0.01] |
| mother-daughter | 0.931 [0.894, 1.000] | 0.260 [0.259, 0.285] | 3.576 [3.230, 4.051] | 0.001 [0.001, 0.002] |

model estimates (mean [minimum, maximum]) for the 10,000 iterations of the model with non-kin as reference level linear mixed model: $\text{sqrt}(\text{grooming duration}) \sim \text{kin class} + (1 + \text{kin class} | \text{group}) + (1 + \text{kin class} | \text{year}) + (1 | \text{dyad}) + (1 | \text{focal\_animal}) + (1 | \text{action\_partner})$ t score & p value for intercept not shown because of limited interpretation

**Figure 1:**
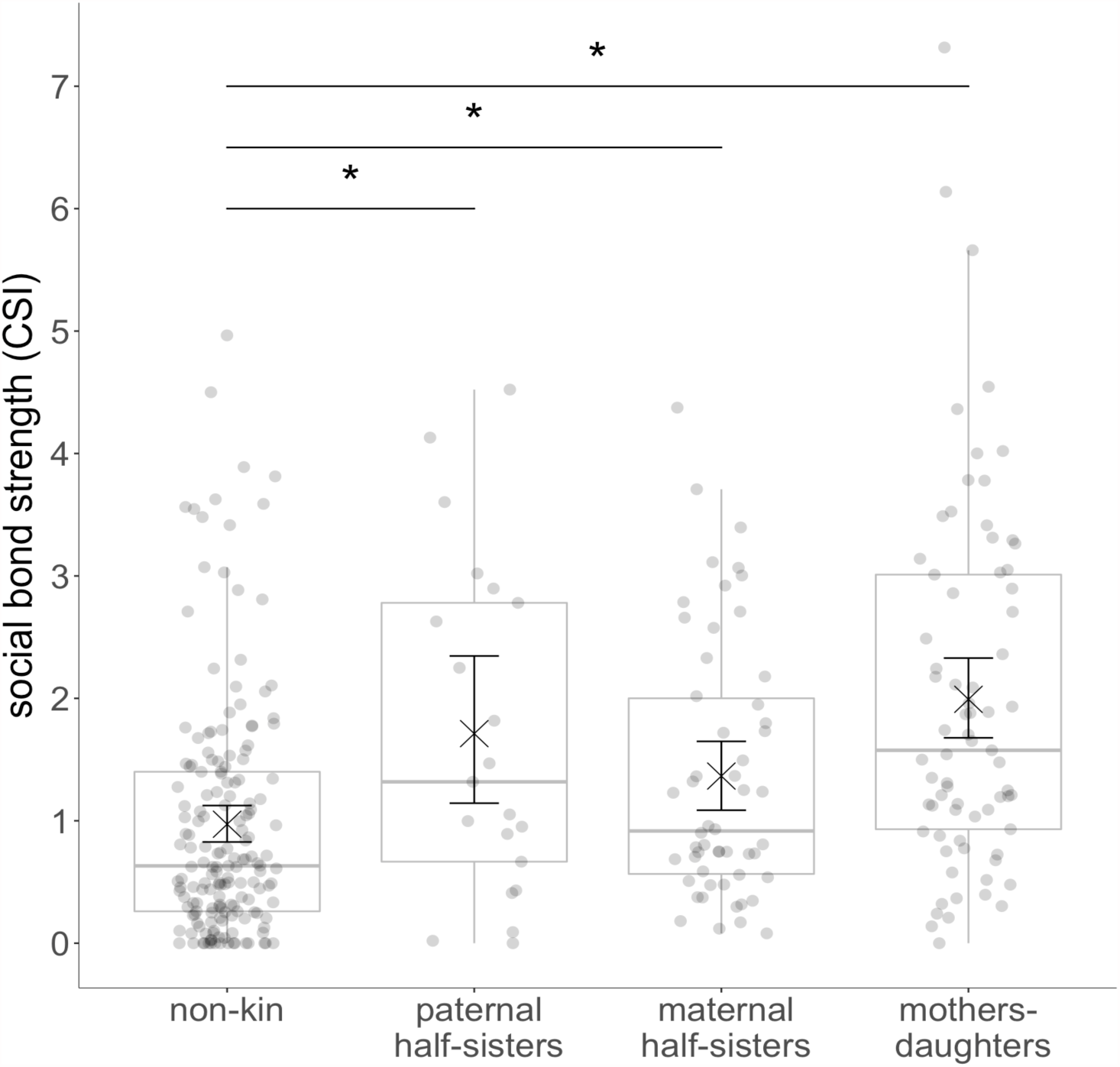
Effect of kin class on dyadic social bond strength. Points depict dyads of adult females in each kin class. Boxplots depict the median, the lower and upper quartiles (25% and 75%), and the range excluding outliers. (x) depict the mean, and the whiskers around it depict the 95% confidence intervals.

**Figure 2:**
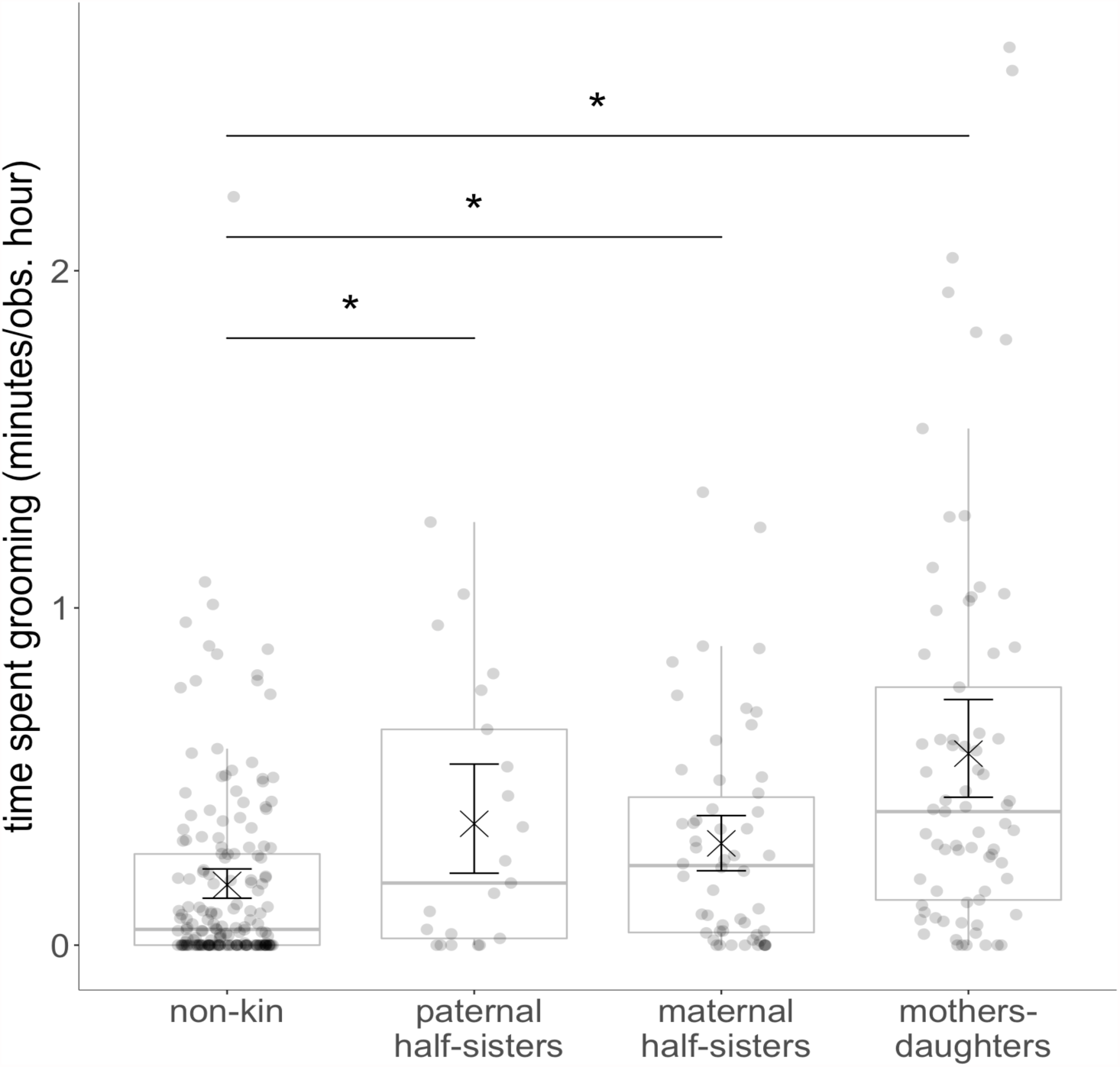
Effect of kin class on time spent grooming. Points depict dyads of adult females in each kin class. Boxplots depict the median, the lower and upper quartiles (25% and 75%), and the range excluding outliers. (x) depict the mean, and the whiskers around it depict the 95% confidence intervals.

### Effect of close maternal kin availability on paternal kin biases (Model 3)

The full model with kin class and sum of close maternal kin available (mothers, adult daughters and adult maternal half-sisters) as predictors of bond strength was a significantly better fit to the data than the null model (likelihood ratio test: χ^2^ = 8.8 [4.5, 11.3], df = 2, p = 0.01 [0.003, 0.1], mean [minimum, maximum] over 10,000 tests). Because the interaction between kin class and sum of close maternal kin available was not significant, the interaction term was excluded in the final model. Together, kin class and sum of close maternal kin available explained some of the variation in the data (marginal R^2^= 0.05 [0.03, 0.07], mean [minimum, maximum] over 10,000 tests). As in the previous models, paternal half-sisters formed significantly stronger bonds than the non-kin dyads. The sum of close maternal kin available had no significant effect on social bond strength, but especially for the paternal half-sisters the sample size might be too small to detect such an effect (see Figure 3 & Table 5). Moreover, this finding should be interpreted with caution, as there were most likely more maternal kin available in the dyads we could not reliably assign to a kin class.

**Table 5:** Effect of maternal kin availability on dyadic social bond strength of paternal half-sisters and non-kin

| Term | estimate | Standard Error (SE) | t score | p value |
| --- | --- | --- | --- | --- |
| Intercept | 0.839 [0.822, 0.862] | 0.045 [0.044, 0.063] | - | - |
| paternal half-sisters | 0.360 [0.265, 0.424] | 0.139 [0.131, 0.141] | 2.599 [1.928, 3.099] | 0.03 [0.03, 0.03] |
| sum of close maternal kin | 0.071 [0.003, 0.104] | 0.042 [0.042, 0.048] | 1.679 [0.067, 2.375] | 0.22 [0.22, 0.22] |

model estimates (mean [minimum, maximum]) for the 10,000 iterations of the model with non-kin as reference level linear mixed model: $\text{sqrt(CSI)} \sim \text{kin class} + \text{z.sum of close maternal kin} + (1 + \text{z.sum close maternal kin} | \text{group}) + (1 + \text{z.sum close maternal kin} | \text{year}) + (1 | \text{dyad}) + (1 | \text{focal\_animal}) + (1 | \text{action\_partner})$ t score & p value for intercept not shown because of limited interpretation

**Figure 3:**
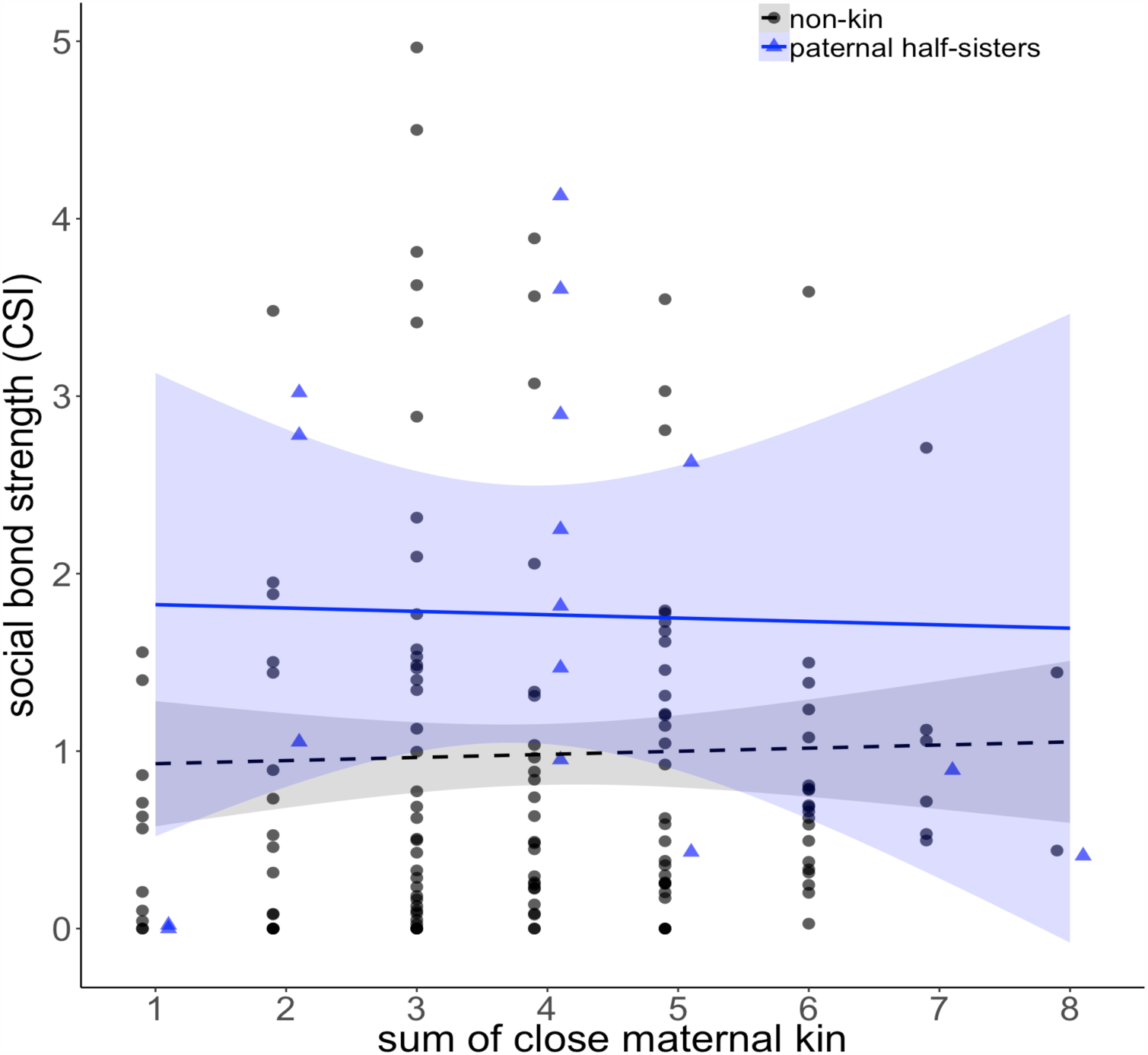
Effect of maternal kin availability on dyadic social bond strength of paternal half-sisters and non-kin. Black dots depict non-kin dyads, with the dotted black line depicting the least squares linear regression line with the 95% confidence interval around it. Blue triangles depict paternal half-sister dyads with the solid blue line depicting the least squares linear regression line with the 95% confidence interval around it.

### Effect of age difference on paternal kin biases (Model 4&5)

The full model with kin class, age difference, and their interaction as predictors of bond strength fitted the data significantly better than the null model (likelihood ratio test: χ^2^ = 6.5 [4.4, 8.8], df = 2, p = 0.04 [0.01, 0.11], mean [minimum, maximum] over 10,000 tests). The interaction between kin class and age difference was not significant, so interaction term was removed from the final model. Together, kin class and age difference explained some of the variation in social bond strength (marginal R^2^= 0.03 [0.03, 0.05], mean [minimum, maximum] over 10,000 tests). As in the previous model with the full dataset, bonds between paternal half-sisters were significantly stronger than bonds between non-kin dyads. Age difference had no significant effect on social bond strength (Figure 4 & Table 6).

**Table 6:** Effect of age difference on dyadic social bond strength of paternal half-sisters and non-kin

| Term | estimate | Standard Error (SE) | t score | p value |
| --- | --- | --- | --- | --- |
| Intercept | 0.848 [0.833, 0.866] | 0.041 [0.039, 0.057] | - | - |
| paternal half-sisters | 0.310 [0.209, 0.359] | 0.119 [0.113, 0.121] | 2.614 [1.781, 3.128] | 0.02 [0.02, 0.02] |
| age difference | -0.046 [-0.070, -0.029] | 0.037 [0.037, 0.041] | -1.225 [-1.805, -0.725] | 0.52 [0.52, 0.52] |

model estimates (mean [minimum, maximum]) for the 10,000 iterations of the model with non-kin as reference level linear mixed model: $\text{sqrt(CSI)} \sim \text{kin class} + \text{sqrt(z.age difference)} + (1|\text{dyad}) + (1|\text{focal\_animal}) + (1|\text{action\_partner})$ t score & p value for intercept not shown because of limited interpretation

**Figure 4:**
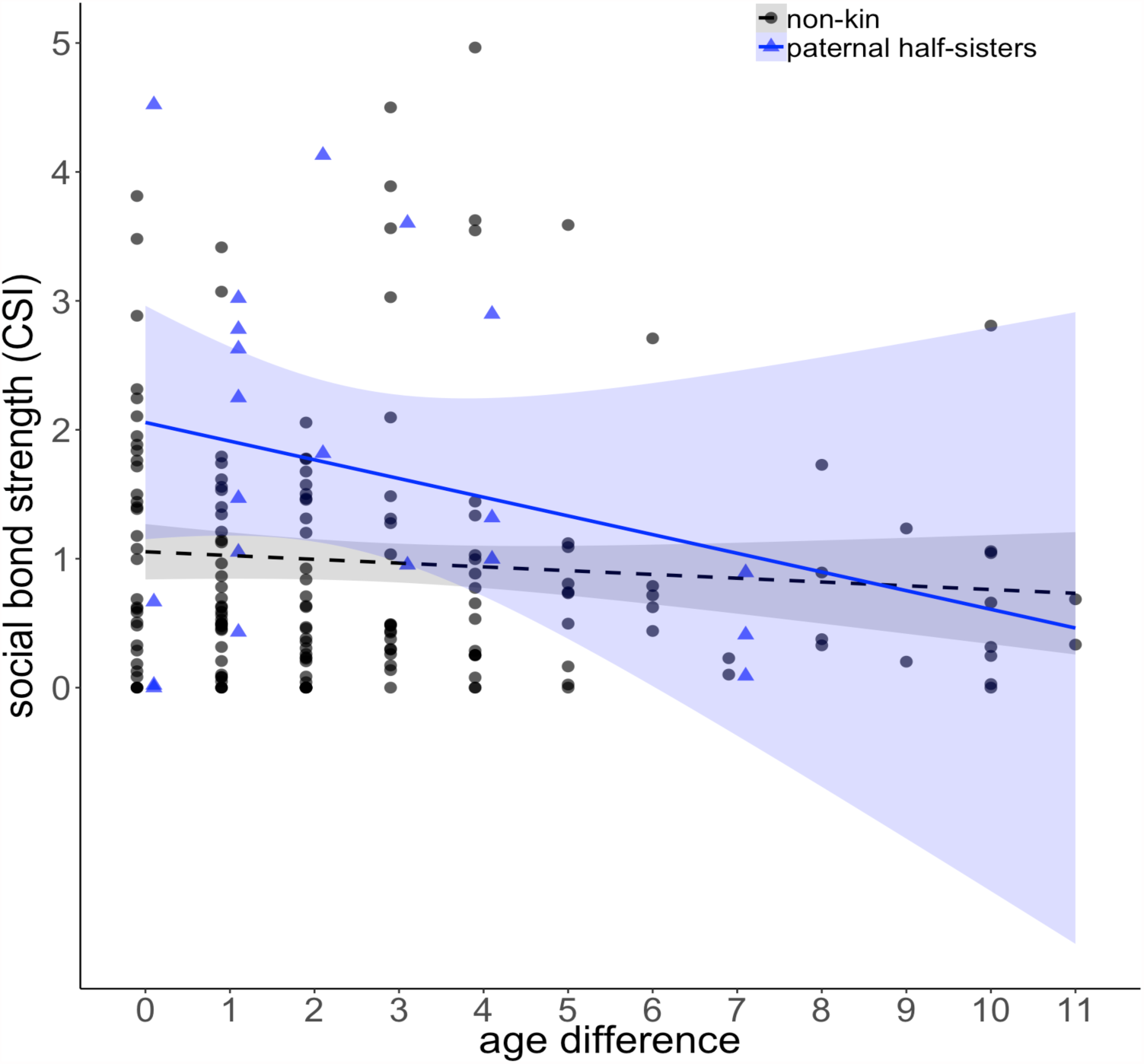
Effect of age difference on dyadic social bond strength of paternal half-sisters and non-kin. Black dots depict non-kin dyads, with the dotted black line depicting the least squares linear regression line with the 95% confidence interval around it. Blue triangles depict paternal half-sister dyads with the solid blue line depicting the least squares linear regression line with the 95% confidence interval around it.

When running the model with a binary variable coding for whether partners were born the same year or not, the full model was a better fit to the data than the null model too [likelihood ratio test: χ^2^ = 8.3 [4.3, 12.1], df = 2, p = 0.02 [0.002, 0.12], mean [minimum, maximum] over 10,000 tests). Together kin class and cohort explained some of the variation in social bond strength (marginal R^2^= 0.04 [0.02, 0.06], mean [minimum, maximum] over 10,000 tests). Similar to the age difference model, paternal half-sisters formed significantly stronger bonds than non-kin, but being born in the same year did not impact social bond strength (Figure 5 & Table 7).

**Table 7:** Effect of age cohort on dyadic social bond strength of paternal half-sisters and non-kin

| Term | estimate | Standard Error (SE) | t score | p value |
| --- | --- | --- | --- | --- |
| Intercept | 0.926 [0.899, 0.969] | 0.088 [0.086, 0.094] | - | - |
| paternal half-sisters | 0.315 [0.247, 0.370] | 0.119 [0.113, 0.122] | 2.660 [2.118, 3.209] | 0.018 [0.016, 0.020] |
| age cohort | -0.095 [-0.154, -0.064] | 0.096 [0.094, 0.100] | -0.998 [-1.613, -0.668] | 0.611 [0.611, 0.611] |

model estimates (mean [minimum, maximum]) for the 10,000 iterations of the model with non-kin as reference level linear mixed model: $\text{sqrt(CSI)} \sim \text{kin class} + \text{sqrt(age cohort)} + (1|\text{dyad}) + (1|\text{focal\_animal}) + (1|\text{action\_partner})$ t score & p value for intercept not shown because of limited interpretation

**Figure 5:**
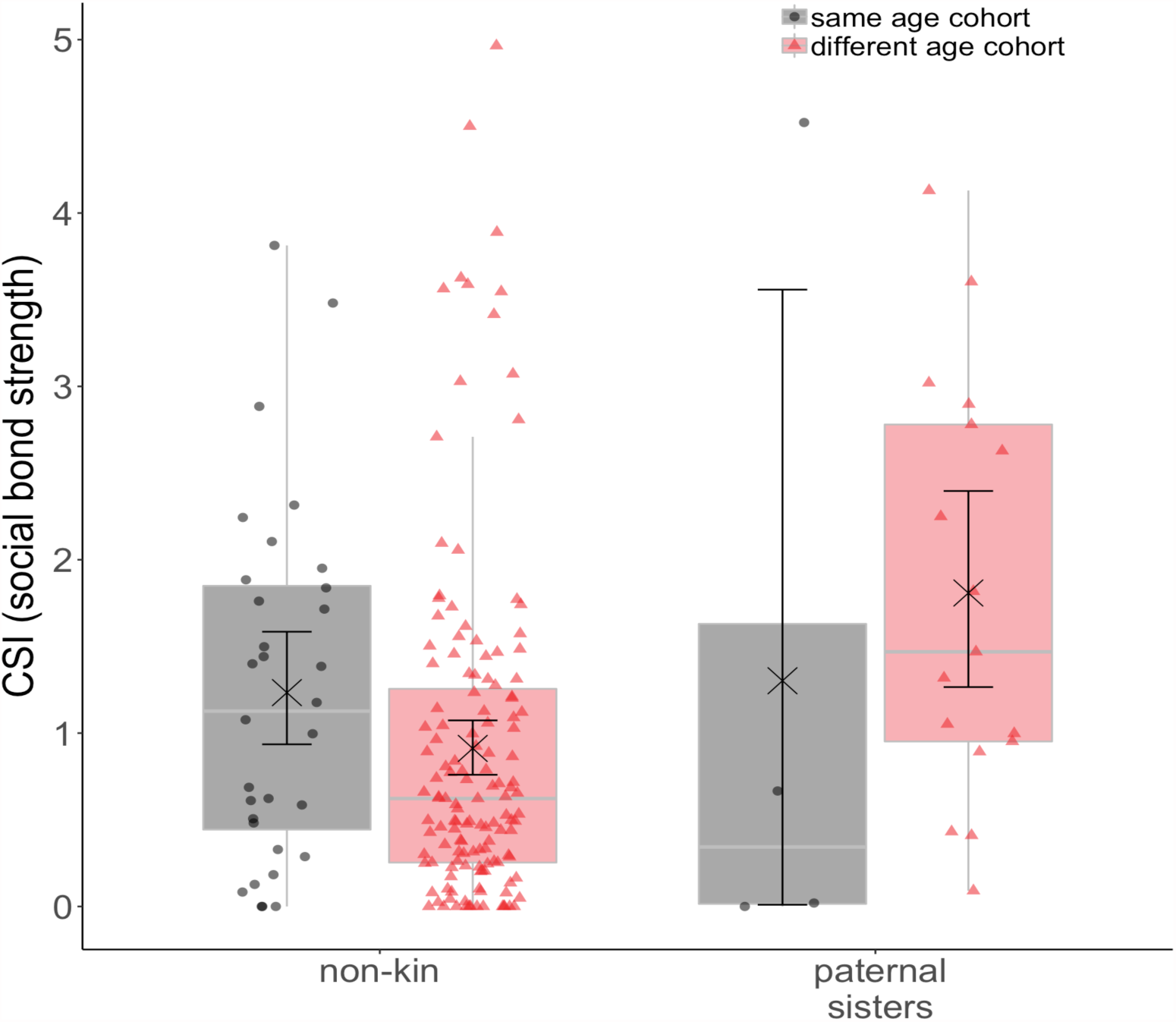
Effect of age cohort on dyadic social bond strength of paternal half-sisters and non-kin. Black dots and boxplots depict dyads born in the same age cohort, red triangles and boxplots depicts dyads born in different age cohorts. Boxplots depict the median, the lower and upper quartiles (25% and 75%), and the range excluding outliers. (x) depict the mean, and the whiskers around it depict the 95% confidence intervals.

### Kin effects in group fission

Another source of evidence for the importance of maternal kinship in our species comes from a group fission that occurred in one of our study groups in 2012 and led to a permanent fission into two subgroups. During this event, two entire matrilines split off from the original group, leaving no maternally related females behind and taking no other female with them.

## DISCUSSION

Kin biases are expected to develop if kin are available and reliably recognisable. In most mammal species, maternal kin meet both criteria, and maternal relatedness has emerged as one of the most important factors structuring mammal social organization and behaviour (Silk, 2009; J. E. Smith, 2014). Especially in stable social groups with female philopatry and overlap of generations, strong mother-offspring bonds lead to the clustering of females into matrilines, with maternal kin being closely associated and selectively directing affiliation towards each other (Archie et al., 2006; Berman, 2015; J. E. Smith et al., 2010). As expected, we found that maternal kinship impacted the sociality of female Assamese macaques. Maternal half-sisters and mother-daughter dyads both formed stronger social bonds and spent more time grooming than non-kin. The importance of maternal kinship was also reflected in the group fission we observed, which happened fully along matrilines. This is in line with several studies in other mammal species that find that both temporal fission in species with fission-fusion dynamics as well as permanent group splits are predicted by genetic relationships, and especially maternal relatedness (Archie et al., 2006; Carter, Seddon, Frère, Carter, & Goldizen, 2013; Van Horn, Buchan, Altmann, & Alberts, 2007).

In contrast to maternal kin, paternal kin availability and recognition are less well understood, and depend on species-specific factors such as dispersal and mating regime (Strier, 2004). Since most mammal females mate with multiple partners during their fertile period, paternity is concealed and individuals rely on proxies for paternal kin recognition (Widdig, 2007). Whether animals can assess paternal relatedness from such proxies, and how reliable those cues are is currently under debate (Langergraber, 2012; Widdig, 2007, 2013). To date, few studies have looked into how paternal relatedness affects sociality (Table 8), making it hard to draw conclusions on which factors affect the development of biases towards paternal kin.

**Table 8:**
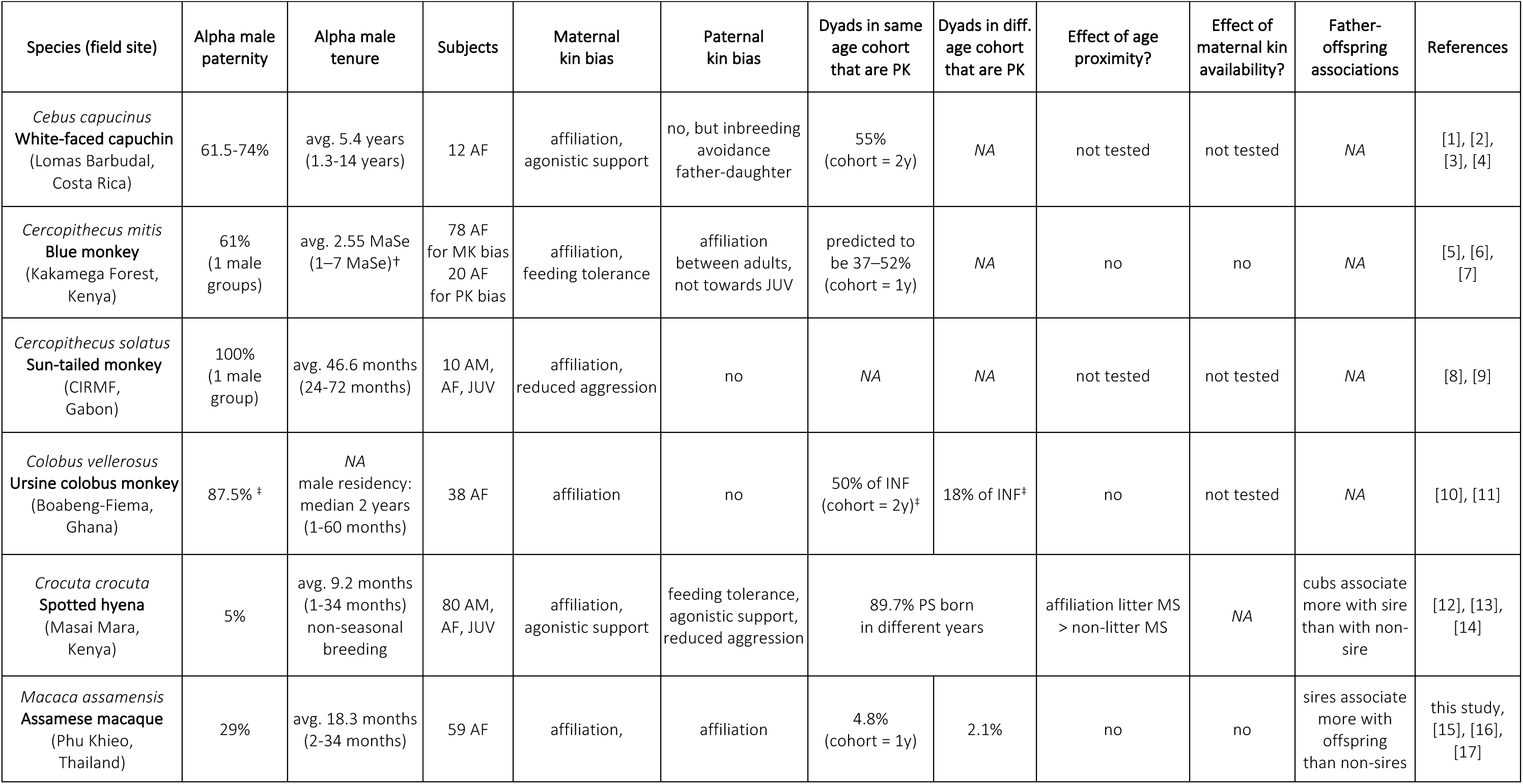
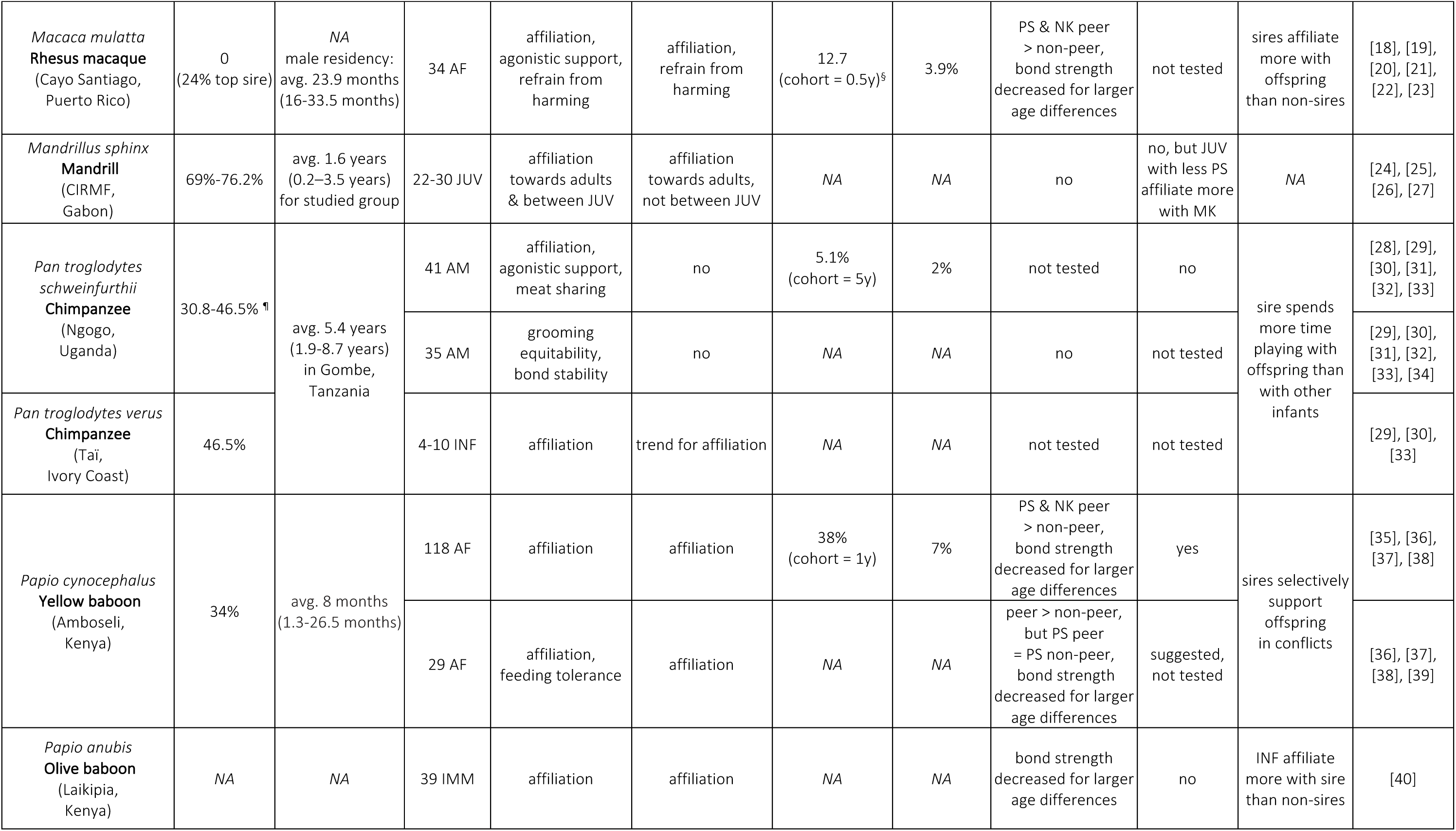
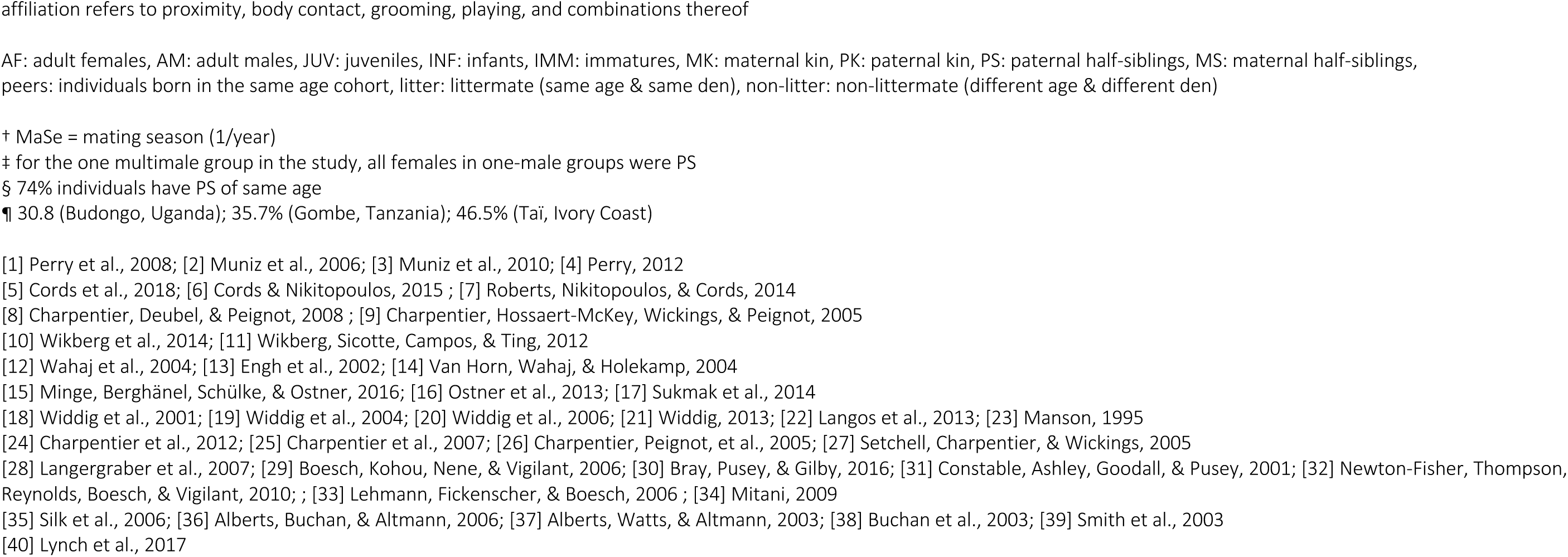
Demographic and mating patterns of species studied on paternal kin biases

| Species (field site) | Alpha male paternity | Alpha male tenure | Subjects | Maternal kin bias | Paternal kin bias | Dyads in same age cohort that are PK | Dyads in diff. age cohort that are PK | Effect of age proximity? | Effect of maternal kin availability? | Father-offspring associations | References |
| --- | --- | --- | --- | --- | --- | --- | --- | --- | --- | --- | --- |
| *Cebus capucinus*<br>**White-faced capuchin**<br>(Lomas Barbudal, Costa Rica) | 61.5-74% | avg. 5.4 years<br>(1.3-14 years) | 12 AF | affiliation,<br>agonistic support | no, but inbreeding avoidance<br>father-daughter | 55%<br>(cohort = 2y) | NA | not tested | not tested | NA | [1], [2],<br>[3], [4] |
| *Cercopithecus mitis*<br>**Blue monkey**<br>(Kakamega Forest, Kenya) | 61%<br>(1 male groups) | avg. 2.55 MaSe<br>(1-7 MaSe)<sup>†</sup> | 78 AF<br>for MK bias<br>20 AF<br>for PK bias | affiliation,<br>feeding tolerance | affiliation<br>between adults,<br>not towards JUV | predicted to be 37-52%<br>(cohort = 1y) | NA | no | no | NA | [5], [6],<br>[7] |
| *Cercopithecus solatus*<br>**Sun-tailed monkey**<br>(CIRMF, Gabon) | 100%<br>(1 male group) | avg. 46.6 months<br>(24-72 months) | 10 AM,<br>AF, JUV | affiliation,<br>reduced aggression | no | NA | NA | not tested | not tested | NA | [8], [9] |
| *Colobus vellerosus*<br>**Ursine colobus monkey**<br>(Boabeng-Fiema, Ghana) | 87.5%<sup>‡</sup> | NA<br>male residency:<br>median 2 years<br>(1-60 months) | 38 AF | affiliation | no | 50% of INF<br>(cohort = 2y)<sup>‡</sup> | 18% of INF<sup>‡</sup> | no | not tested | NA | [10], [11] |
| *Crocuta crocuta*<br>**Spotted hyena**<br>(Masai Mara, Kenya) | 5% | avg. 9.2 months<br>(1-34 months)<br>non-seasonal<br>breeding | 80 AM,<br>AF, JUV | affiliation,<br>agonistic support | feeding tolerance,<br>agonistic support,<br>reduced aggression | 89.7% PS born<br>in different years |  | affiliation litter MS<br>> non-litter MS | NA | cubs associate<br>more with sire<br>than with non-sire | [12], [13],<br>[14] |
| *Macaca assamensis*<br>**Assamese macaque**<br>(Phu Khieo, Thailand) | 29% | avg. 18.3 months<br>(2-34 months) | 59 AF | affiliation, | affiliation | 4.8%<br>(cohort = 1y) | 2.1% | no | no | sires associate<br>more with<br>offspring<br>than non-sires | this study,<br>[15], [16],<br>[17] |
| *Macaca mulatta*<br>**Rhesus macaque**<br>(Cayo Santiago,<br>Puerto Rico) | 0<br>(24% top sire) | NA<br>male residency:<br>avg. 23.9 months<br>(16-33.5 months) | 34 AF | affiliation,<br>agonistic support,<br>refrain from<br>harming | affiliation,<br>refrain from<br>harming | 12.7<br>(cohort = 0.5y)<sup>§</sup> | 3.9% | PS & NK peer<br>> non-peer,<br>bond strength<br>decreased for larger<br>age differences | not tested | sires affiliate<br>more with<br>offspring<br>than non-sires | [18], [19],<br>[20], [21],<br>[22], [23] |
| *Mandrillus sphinx*<br>**Mandrill**<br>(CIRMF,<br>Gabon) | 69%-76.2% | avg. 1.6 years<br>(0.2–3.5 years)<br>for studied group | 22-30 JUV | affiliation<br>towards adults<br>& between JUV | affiliation<br>towards adults,<br>not between JUV | NA | NA | no | no, but JUV<br>with less PS<br>affiliate more<br>with MK | NA | [24], [25],<br>[26], [27] |
| *Pan troglodytes*<br>*schweinfurthii*<br>**Chimpanzee**<br>(Ngogo,<br>Uganda) | 30.8-46.5%<sup>¶</sup> | avg. 5.4 years<br>(1.9-8.7 years)<br>in Gombe,<br>Tanzania | 41 AM | affiliation,<br>agonistic support,<br>meat sharing | no | 5.1%<br>(cohort = 5y) | 2% | not tested | no | sire spends<br>more time<br>playing with<br>offspring than<br>with other<br>infants | [28], [29],<br>[30], [31],<br>[32], [33] |
|  |  |  | 35 AM | grooming<br>equitability,<br>bond stability | no | NA | NA | no | not tested |  | [29], [30],<br>[31], [32],<br>[33], [34] |
| *Pan troglodytes verus*<br>**Chimpanzee**<br>(Taï,<br>Ivory Coast) | 46.5% |  | 4-10 INF | affiliation | trend for affiliation | NA | NA | not tested | not tested |  | [29], [30],<br>[33] |
| *Papio cynocephalus*<br>**Yellow baboon**<br>(Amboseli,<br>Kenya) | 34% | avg. 8 months<br>(1.3-26.5 months) | 118 AF | affiliation | affiliation | 38%<br>(cohort = 1y) | 7% | PS & NK peer<br>> non-peer,<br>bond strength<br>decreased for larger<br>age differences | yes | sires selectively<br>support<br>offspring<br>in conflicts | [35], [36],<br>[37], [38] |
|  |  |  | 29 AF | affiliation,<br>feeding tolerance | affiliation | NA | NA | peer > non-peer,<br>but PS peer<br>= PS non-peer,<br>bond strength<br>decreased for larger<br>age differences | suggested,<br>not tested |  | [36], [37],<br>[38], [39] |
| *Papio anubis*<br>**Olive baboon**<br>(Laikipia,<br>Kenya) | NA | NA | 39 IMM | affiliation | affiliation | NA | NA | bond strength<br>decreased for larger<br>age differences | no | INF affiliate<br>more with sire<br>than non-sires | [40] |

affiliation refers to proximity, body contact, grooming, playing, and combinations thereof
AF: adult females, AM: adult males, JUV: juveniles, INF: infants, IMM: immatures, MK: maternal kin, PK: paternal kin, PS: paternal half-siblings, MS: maternal half-siblings, peers: individuals born in the same age cohort, litter: littermate (same age & same den), non-litter: non-littermate (different age & different den)
† MaSe = mating season (1/year) ‡ for the one multimale group in the study, all females in one-male groups were PS § 74% individuals have PS of same age ¶ 30.8 (Budongo, Uganda); 35.7% (Gombe, Tanzania); 46.5% (Tai, Ivory Coast)
[1] Perry et al., 2008; [2] Muniz et al., 2006; [3] Muniz et al., 2010; [4] Perry, 2012 [5] Cords et al., 2018; [6] Cords & Nikitopoulos, 2015 ; [7] Roberts, Nikitopoulos, & Cords, 2014 [8] Charpentier, Deubel, & Peignot, 2008 ; [9] Charpentier, Hossaert-McKey, Wickings, & Peignot, 2005 [10] Wikberg et al., 2014; [11] Wikberg, Sicotte, Campos, & Ting, 2012 [12] Wahaj et al., 2004; [13] Engh et al., 2002; [14] Van Horn, Wahaj, & Holekamp, 2004 [15] Minge, Berghänel, Schülke, & Ostner, 2016; [16] Ostner et al., 2013; [17] Sukmak et al., 2014 [18] Widdig et al., 2001; [19] Widdig et al., 2004; [20] Widdig et al., 2006; [21] Widdig, 2013; [22] Langos et al., 2013; [23] Manson, 1995 [24] Charpentier et al., 2012; [25] Charpentier et al., 2007; [26] Charpentier, Peignot, et al., 2005; [27] Setchell, Charpentier, & Wickings, 2005 [28] Langergraber et al., 2007; [29] Boesch, Kohou, Nene, & Vigilant, 2006; [30] Bray, Pusey, & Gilby, 2016; [31] Constable, Ashley, Goodall, & Pusey, 2001; [32] Newton-Fisher, Thompson, Reynolds, Boesch, & Vigilant, 2010; ; [33] Lehmann, Fickenscher, & Boesch, 2006 ; [34] Mitani, 2009 [35] Silk et al., 2006; [36] Alberts, Buchan, & Altmann, 2006; [37] Alberts, Watts, & Altmann, 2003; [38] Buchan et al., 2003; [39] Smith et al., 2003 [40] Lynch et al., 2017

With this study, we addressed some of the gaps in what we know so far on paternal kin biases. Assamese macaques have a relatively low male reproductive skew (29% alpha male paternity: Sukmak et al., 2014), and a relatively long alpha male tenure (Ostner et al., 2013), which allowed us to test whether an affiliation bias towards paternal kin can develop in a species where several individuals are expected to have paternal half-siblings available, but where those paternal half-sisters usually belong to different age cohorts. Out of the 59 observed adult females, 18 females had at least one paternal half-sister in the group for at least one of the years of our study. Only 2 of the 12 (12.5%) paternal half-sister dyads were born in the same year (average age difference 2.7a). Two of the 42 (4.8%) dyads born in the same year were paternal half-sisters, compared to 10 of 471 dyads (2.1%) born in different years.

We found that female Assamese macaques bias their affiliation towards paternal kin, independent of age similarity. Paternal half-sisters formed stronger bonds and spent more time grooming than did non-kin, but as expected we found no evidence for an effect of age proximity on social bond strength. It seems that familiarity through age proximity is not a prerequisite for paternal kin biases to develop, and that individuals might recognize their paternal kin through other mechanisms. In fact, based on the limited data so far, the importance of familiarity via age proximity is generally questionable (Langergraber, 2012; Widdig, 2013). If individuals discriminate paternal kin solely based on growing up together, one would expect a preference for all individuals (also non-kin) born in the same age cohort, the underlying assumption being that most of those individuals are paternal kin (Altmann, 1979). Paternity is, however, rarely monopolized by one male, and in species in which the alpha male does sire a large proportion of the offspring, he also usually has a relatively long tenure (Widdig, 2013; Table 8). Even in species in which paternal kin biases are more pronounced among age mates, only a small proportion of same-aged individuals are actually paternal kin (Table 8). So, although there is an effect of age proximity in these species, the question is whether this attraction to same-aged individuals really reflects a bias towards paternal kin. It is also important to note that paternal kin biases are more pronounced, but not limited to age mates in all these species (Lynch et al., 2017; K. Smith et al., 2003; Widdig et al., 2001), suggesting that individuals do not rely on age proximity alone to discriminate paternal kin from non-kin. Moreover, similar to our finding, other species show paternal kin biases in affiliation, but no effect of age similarity (Charpentier et al., 2007; Cords et al., 2018). Age proximity thus might serve as a first, rather rough proxy for paternal relatedness in some species, and individuals likely make use of other cues as well to recognize their paternal kin.

An alternative, and so far rather understudied way through which paternal kin could recognize each other is through mother- and/or father-mediated familiarity (Widdig, 2007). In Assamese macaques, for example, males and females form strong, stable bonds that can last for several years (Haunhorst, Schülke, & Ostner, 2016). A male associated with a female during the mating season is her main mating partner, and thus the most likely sire of her offspring (Ostner et al., 2013). In other primate species too, males and females form associations that persist after the female gave birth (Baniel, Cowlishaw, & Huchard, 2016; Huchard et al., 2010; Lemasson, Palombit, & Jubin, 2008; Moscovice et al., 2010; Nguyen, Van Horn, Alberts, & Altmann, 2009; Van Schaik & Kappeler, 1997), which might allow infants to get indirectly familiarized to their father early on in life. Infants might also be directly familiarized to their father through male paternal care (Buchan, Alberts, Silk, & Altmann, 2003; Charpentier, Van Horn, Altmann, & Alberts, 2008; Huchard et al., 2013; Langos, Kulik, Mundry, & Widdig, 2013; Onyango, Gesquiere, Altmann, & Alberts, 2013). Evidence is building for male-infant associations between father and offspring in several species (Table 8; Huchard et al., 2010), which in turn allows for the familiarization of paternally related infants. Alternatively, paternal half-siblings can get familiarized to each other directly if several females share the same preferred male affiliation partner (Haunhorst et al., 2016; Seyfarth & Cheney, 2012). In olive baboon immatures (*Papio anubis*), bonds between paternal half-siblings are stronger when the father is present in the group (Lynch et al., 2017), which highlights the importance of a shared association with a common father for the development of a bias towards paternal kin. In our study, fathers remained in the group for an average of 2.4 years (9 months – 5.7 years) after the birth of two paternal half-sisters, which would have given them ample time to get familiarized to each other through the common father.

Another possibility is that individuals recognize their paternal kin through phenotype matching. Several recent studies have shown that primates and other mammals are capable of using acoustic, olfactory or visual cues to recognize their kin (Charpentier et al., 2017; Gilad, Swaisgood, Owen, & Zhou, 2016; Henkel & Setchell, 2018; Mateo, 2017; Pfefferle, Ruiz-Lambides, & Widdig, 2014, 2015). Phenotype matching and familiarity are not mutually exclusive mechanisms for kin recognition, and animals likely use multiple cues simultaneously to discriminate kin from non-kin (Tang-Martinez, 2001; Widdig, 2007).

Independent of the mechanism behind it, paternal kin recognition is based on proxies more prone to error than the mother-mediated familiarity through which animals likely recognize their maternal kin (Widdig, 2007). It is therefore not surprising that paternal kin biases are usually less pronounced than biases towards maternal kin (Charpentier et al., 2012; Lynch et al., 2017; Silk et al., 2006; Widdig et al., 2001). Contrary to our prediction, however, we found no difference in bond strength between close maternal kin and paternal half-sisters. Three factors might explain why paternal half-sisters formed bonds similar in strength as maternal half-sisters and mother-daughter dyads: (1) paternal kin biases might be modulated by maternal kin availability either if females form close compensatory bonds with their paternal kin because too little close maternal kin is available (Silk et al., 2006; K. Smith et al., 2003), or if females form bonds with paternal kin only if they are not saturated with close maternal kin; (2) paternal kin might be as valuable as bonding partners as maternal kin are and/or (3) we lacked the power to detect a difference in bond strength between maternal and paternal kin (sample size of 59 adult females, 12 paternal half-sister dyads).

We found no evidence for an effect of the number of maternal kin available on the strength of social bonds between paternal half-sisters or non-kin. In fact, the one female in our study that had no (assigned) maternal kin available formed the weakest bond with her paternal half-sister of all bonds between paternal half-sisters. Our results indicate that the close bonds we observed between paternal kin were not compensating for a lack of close maternal kin. It is important to note that to really assess whether females form compensatory bonds with paternal kin would require monitoring how individuals adjust their social network after losing or gaining close maternal kin. In chacma baboons (*Papio ursinus*), females increase the number of grooming partners after the loss of a close relative (Engh et al., 2006), and they might do so by turning towards paternal kin. As we had only one dyad of paternal half-sisters where the number of close maternal kin changed within our study period, we could not look into how females adjust to changes in partner availability in this study.

Paternal kin biases might also only develop if individuals do not saturate their social time on close maternal kin. The fact that we found no effect of maternal kin availability on paternal kin biases might imply that the number of close maternal kin (up to 5 maternal half-sisters, mother and daughters for each female, probably more in the unassigned dyads) is not high enough to saturate the social time available to the females in our study. It might be that maternal kin availability only impacts paternal kin biases in extreme cases, where females have either (almost) no close maternal kin available (K. Smith et al., 2003: similar bias towards paternal and maternal kin), or where too many maternal kin are available (Perry et al., 2008: no paternal kin bias), while in less extreme cases number of close maternal kin available or matriline size do not affect paternal kin biases (this study; Charpentier et al., 2012; Cords et al., 2018).

Since maternal kin availability does not explain why we found similar biases towards paternal and maternal kin, paternal kin could in fact be preferred to the same extent as maternal kin. The social behaviours that we studied (proximity, body contact and grooming) are not costly behaviours. Even if paternal kin are recognized less accurately than maternal kin, the cost of investing any of those behaviours in a wrongly discriminated non-related individual would be very small. Biases towards paternal kin in relatively low-cost behaviours might thus be more pronounced than in costlier behaviours such as agonistic support. Rhesus macaques for example refrain from harming their paternal kin, rather than actively supporting them (Widdig et al., 2006). Additionally, paternal kin might be valuable bonding partners, especially for lower-ranking females, as they might have larger rank differences, in contrast to maternal kin who are usually close in rank (Lynch et al., 2017). Social bonds in female Assamese macaques enhance feeding tolerance, so bonding with higher-ranking paternal half-sister might increase access to valuable resources (Heesen, Rogahn, Macdonald, Ostner, & Schülke, 2014). In mandrills (*Mandrillus sphinx*), juveniles form stronger bonds with their distant maternal kin when they have less paternal kin available (Charpentier et al., 2012). In olive baboons, immatures who had a father present in the group formed weaker bonds with their maternal half-siblings (Lynch et al., 2017). It might thus be that animals have some flexibility in the choice of their bonding partners, and that when they are able to discriminate paternal kin with some reliability, they might be as valuable partners as maternal kin.

On a broader level, our data contributes in differentiating between two perspectives on how reproductive skew drives relatedness patterns, and how relatedness in turn drives social structure. The first perspective is that in species with a high reproductive skew, most group members are closely related. Selection will then favour indifferent tolerance towards all group members for inclusive fitness benefits (Lukas & Clutton-Brock, 2018). The second perspective is that tolerance is reserved for kin only. In species with low male reproductive skew and short male tenures few paternal kin are present in the group and all kin is maternal, yielding networks of largely isolated matrilineal clusters. In contrast, in species with high paternity concentration and/or long reproductive tenures many paternal kin might be present. Kin biases towards paternal kin might form bridges of tolerance between matrilines that increase the overall tolerance and break up the substructure at the group level (Schülke & Ostner, 2008). The crucial difference between both perspectives is whether tolerance is differentiated or not. Here we show that Assamese macaque females direct their affiliation selectively towards both paternal and maternal kin over non-kin, which is in line with the second perspective.

## ACKNOWLEDGEMENTS

We thank the National Research Council of Thailand and the Department of National Parks, Wildlife and Plant Conservation for permission to conduct this study and for all the support granted. We are grateful to J. Prabnasuk, K. Nitaya T. Wonsnak, M. Pongjantarasatien and K. Kreetiyutanont, M. Kumsuk, W. Saenphala from Phu Khieo Wildlife Sanctuary for their cooperation over the years and permission to carry out this study. We thank A. Koenig and C. Borries, who established the field site. Special thanks go to S. Jumrudwong, W. Nueorngshiyos, N. Juntuch, J. Wanart, R. Intalo, T. Kilawit, N. Pongangan, B. Klaewklar, N. Bualeng, A. Ebenau, P. Saisawatdikul, K. Srithorn, M. Swagemakers and T. Wisate for their excellent help in the field. We also thank C. Schwarz for her support in the genetics lab, R. Mundry for valuable statistical advice, A.V. Rincon for help in using R, and G. Dezecache for useful comments on the manuscript. We thank the members of the research training group ‘Understanding Social Relationships’ (RTG 2070) for stimulating discussions. This research was funded by the Deutsche Forschungsgemeinschaft (DFG, German Research Foundation) – project number 453 254142454 / GRK 2070.

## AUTHOR CONTRIBUTIONS

J.O. and O.S. designed the research. D.D.M. performed the field research and the lab work, and did the data analyses. J.O. and O.S. supervised the behavioural data collection and analysis, and C.R. advised and supervised the lab work. All authors discussed the results, and D.D.M. took the lead in writing the manuscript. All authors provided critical feedback and helped shape the research, analysis and manuscript.

## DATA ACCESSIBILITY

In case of acceptance data will made accessible at a public repository (Dryad).

